# Nanomechanics of negatively supercoiled diaminopurine-substituted DNA

**DOI:** 10.1101/2021.02.24.432629

**Authors:** Domenico Salerno, Francesco Mantegazza, Valeria Cassina, Matteo Cristofalo, Qing Shao, Laura Finzi, David Dunlap

## Abstract

Single molecule experiments have demonstrated a progressive transition from a B- to an L-form helix as DNA is gently stretched and progressively unwound. Since the particular sequence of a DNA segment influences both base stacking and hydrogen bonding, the conformational dynamics of B-to-L transitions should be tunable. To test this idea, DNA with diaminopurine replacing adenine was synthesized to produce linear fragments with triply hydrogen-bonded A:T base pairs. Triple hydrogen bonding stiffened the DNA by 30% flexurally. In addition, DAP-substituted DNA formed plectonemes with larger gyres for both B- and L-form helices. Both unmodified and DAP-substituted DNA transitioned from a B- to an L-helix under physiological conditions of mild tension and unwinding. This transition avoids writhing by DNA stretched and unwound by enzymatic activity. The intramolecular nature and ease of this transition likely prevent cumbersome topological rearrangements in genomic DNA that would require topoisomerase activity to resolve. L-DNA displayed about tenfold lower persistence length indicating it is much more contractile and prone to sharp bends and kinks. However, left-handed DAP DNA was twice as stiff as unmodified L-DNA. Thus, significantly doubly and triply hydrogen bonded segments have very distinct mechanical dynamics at physiological levels of negative supercoiling and tension.

## Introduction

The conformation and flexibility of DNA depend on electrostatics, base pairing, and base stacking. These interactions produce sequence specific characteristics that influence topology and protein binding. Denaturation directly reflects the hybridization of DNA but does not reveal manifold conformational dynamics of the base-paired state, which confront enzymes, that package, untangle, and process the information contained in DNA. In addition, such enzymes must contend with a continuum of flexural and torsional DNA rigidity conferred by oxidative damage or natural modifications like methylation. Other structural changes can be produced chemically to modify the electrostatic, aromatic, or hydrogen bonding of base pairs (1). For example, 2, 6-diaminopurine (DAP) is an alternative nucleobase that can substitute adenine in A:T base pairs to add an exocyclic amine moiety in the minor groove (2). This amine group forms an additional hydrogen bond with the C2 carbon of thymine in a DAP:T base pair. Complete substitution of adenine-with DAP-deoxyribonucleotide triphosphate in polymerase chain reactions produces triple hydrogen bonding throughout the amplicon. While this modification seems quite uncommon in nature (3,4), it could be useful in DNA-based nanomachines or origami. Such constructs can also serve to probe the activity of enzymes that modify DNA topology or process genetic information (5), since the substitution of DAP for adenine changes base stacking and hydrogen bonding without altering the charge along the molecule.

Substitution of DAP for adenine is known to increase the melting temperature of the B-form (1,6,7). Sequence-specific effects persist and the Santa Lucia model for calculating the melting temperature (8) can be adjusted by uniformly scaling the dinucleotide enthalpies (9). Other data regarding the cyclization of DAP-substituted DNA (DAP-DNA) fragments or binding and obligate tight curling around histone octamers to form nucleosomes indicates that DAP-DNA is flexurally stiffer than unmodified DNA (10,11,12). This is substantiated by increased stiffness exhibited by DAP-DNA molecules a few hundred-to kilo-bases in length deposited on surfaces for atomic force microscopy (9,11,12) and the increased persistence lengths determined using optical or magnetic tweezers to stretch single molecules of DAP-DNA (9,12). Overtwisting DAP-DNA under slight tension also exhibits increased flexural stiffness associated with creating plectonemic coils (5). While the persistence length of DAP-DNA reflects increased rigidity, the overstretching transition of this DNA occurs at a lower tension (9). Thus, DAP-DNA is more rigid and also more susceptible to mechanically induced, structural, phase changes.

Perhaps the most striking phase change exhibited by DAP-DNA is the conversion from a right- to a left-handed helical form. Although DAP-substitution maintains the charge density, the molecule adopts an unusual X-DNA conformation in response to slight concentrations of magnesium in potassium phosphate buffer or 50% concentrations of ethanol (13). This has been shown to be a left-handed helical form with a zigzag phosphate backbone due to alternating syn- and anti-deoxyribose orientations (14). Circular dichroism studies of oligonucleotides of dGC repeats show that high salt concentrations, approximately 60% solutions of alcohols, or added nickel drive the formation of left-handed Z-DNA (14, 15). Critical to this conformational change is the dehydration of the minor groove. Substituting diaminopurine for adenine has a similar effect by inserting an exocyclic amine that displaces water from the minor groove, but in an appropriate solvent stabilizes the X-DNA form, which displays a highly negative CD signal at 280 nm (13). The fact that X-form DAP-DNA is recognized by Z-DNA antibodies and P^31^NMR studies show phosphate signals similar to those recorded for poly(dAT) in an X-DNA form indicate a zigzag backbone.

It is noteworthy that low levels of unwinding and tension have also been shown to drive the transition from right- to left-handed helices in GC segments of DNA (16), which suggested that triple hydrogen bonding may favor the right- to-left transition. To investigate this possibility, magnetic tweezers were used to gently stretch and unwind both unmodified and DAP-substituted DNA. Magnetic tweezers area well established single molecule technique for the simultaneous application of torsion and tension to a single DNA molecule (17,18,19,20,21). By studying resulting variations of the extension of the molecule, we can explore the consequence of DAP substitution on nanomechanical characteristics of DNA filament in the different phases resulting in conditions of high negative torsion.

Overall, we report that the transition to a left-handed form occurred at low levels of unwinding and tension, and left-handed DAP DNA was significantly stiffer than unmodified L-DNA. The tenfold lower persistence length of this left-handed form and the higher susceptibility of triply hydrogen bonded regions to undergo this transition greatly expands the conformational complexity of DNA.

## MATERIAL AND METHODS

### Preparation of DAP and WT DNA

WT and DAP DNA were prepared as previously described (9). Briefly, for the MT experiments T7 DNA ligase was used to ligate a ∼1.0 kbp, multiply digoxigenin-labeled (dig-tail) DNA fragment at one end of a 4642 bp (main) DNA fragment and a ∼0.9 kbp, multiply biotin-labeled (bio-tail) DNA fragment to the other end.

### Magnetic tweezers setup and measurements

We used a custom made Magnetic Tweezers (MT) setup previously described in (22,23,24,25), consisting of an inverted optical microscope equipped with an oil-immersion objective (NIKON 100x, NA = 1.25) mounted on a piezoelectric focusing system (PIFoc, Physik Instrumente, Bresso, Italy). The objective, coupled with a 15 [cm] focal-length lens, led to a 75x magnification. The magnetic field was generated by two permanent neodymium magnets placed above the flow chamber and two piezoelectric motors controlled the position of the magnets along the optical axis (z-direction) and the rotation around the same axis in order to apply a stretching force, or a torque, to the torsionally constrained DNA.

### Microchamber preparation

The flow cell consisted in a square glass capillary tube (1×1 mm^2^ section, 5-cm long, VitroCom, Mountain Lakes, NJ). For each measurement, 250 ml of DNA and magnetic beads suspension were injected, in the absence of a magnetic field, into the capillary previously functionalized as follows. First, a solution of 100 mg/ml polystyrene (average MW230000, Sigma Aldrich) in toluene was injected into the capillary. Then, the capillary was drained and dried with compressed air. In this way, the internal walls were uniformly coated with polystyrene (26). Next, 5 mg of sheep polyclonal anti-digoxigenin antibody (Roche) in 100 ml with 10 mM PBS were incubated in the capillary for 2 h at 37°C. Unbound anti-digoxigenin was eliminated by rinsing the capillary with PBS-Tween-20. The functionalized surface was passivated for 2 h at 37°C with a solution consisting of 10 mM PBS at pH 8 supplemented with 0.1% Tween-20, 1 mg/ml fish sperm DNA (Roche) and 3 mM NaN_3_ (27). Finally, the capillary was rinsed with PBS-Tween-20 and incubated for 1h to allow the DNA to bind to the lower capillary surface. For storage, several capillaries could be prepared simultaneously and kept at 20°C after air drying them.

### Data acquisition and analysis

Images were acquired by a CCD camera (Marlin Allied vision, USA) running at a frame rate of 60 Hz and analyzed in real time by a custom made software developed in Java (Oracle, USA). Calculations on the diffraction rings profile of the magnetic beads allowed to measure the 3D position of the beads with a precision of 10 nm along the optical axes and 40 nm in x-y plane (22). The extracted data was recorded on the pc drive for successive analysis that was performed with an ad hoc software written in MatLab (Mathwork inc. USA).

## RESULTS

### Phenomenology of the torsional behavior of WT and DAP DNA

In this work, we studied the difference between the nanomechanical behavior of WT and DAP DNA in conditions of high negative torsion. Representative twist experiments are reported in Figure 1A where the DNA extension L_e_ is measured as a function of the imposed turns, n_t_, or the corresponding supercoiling, *σ*=n_t_/(N_b_/10.4), where N_b_=4642 is the number of base pairs of the DNA construct. Figure 1A shows data acquired over a large range of imposed turns (from n_t_=+100 until n_t_=-900 turns) for WT and DAP DNA at three different forces F (0.2, 1.1, 2.3 [pN]). Additional data at different forces are reported in Figure S1. Whereas at low forces (F=0.2 [pN]), the behavior of WT and DAP DNA was basically indistinguishable, at intermediate (F=1.1 [pN]) and high (F=2.3 [pN]) forces, the two types of DNA show L_e_ vs n_t_ curves having qualitatively and quantitatively different characteristics. In particular, at intermediate and high forces the extension unwound DNA appeared systematically higher for DAP DNA than for WT DNA. Furthermore, for high forces (F=2.3 [pN]) three different linear regimes are clearly distinguishable.

**Figure 1.**
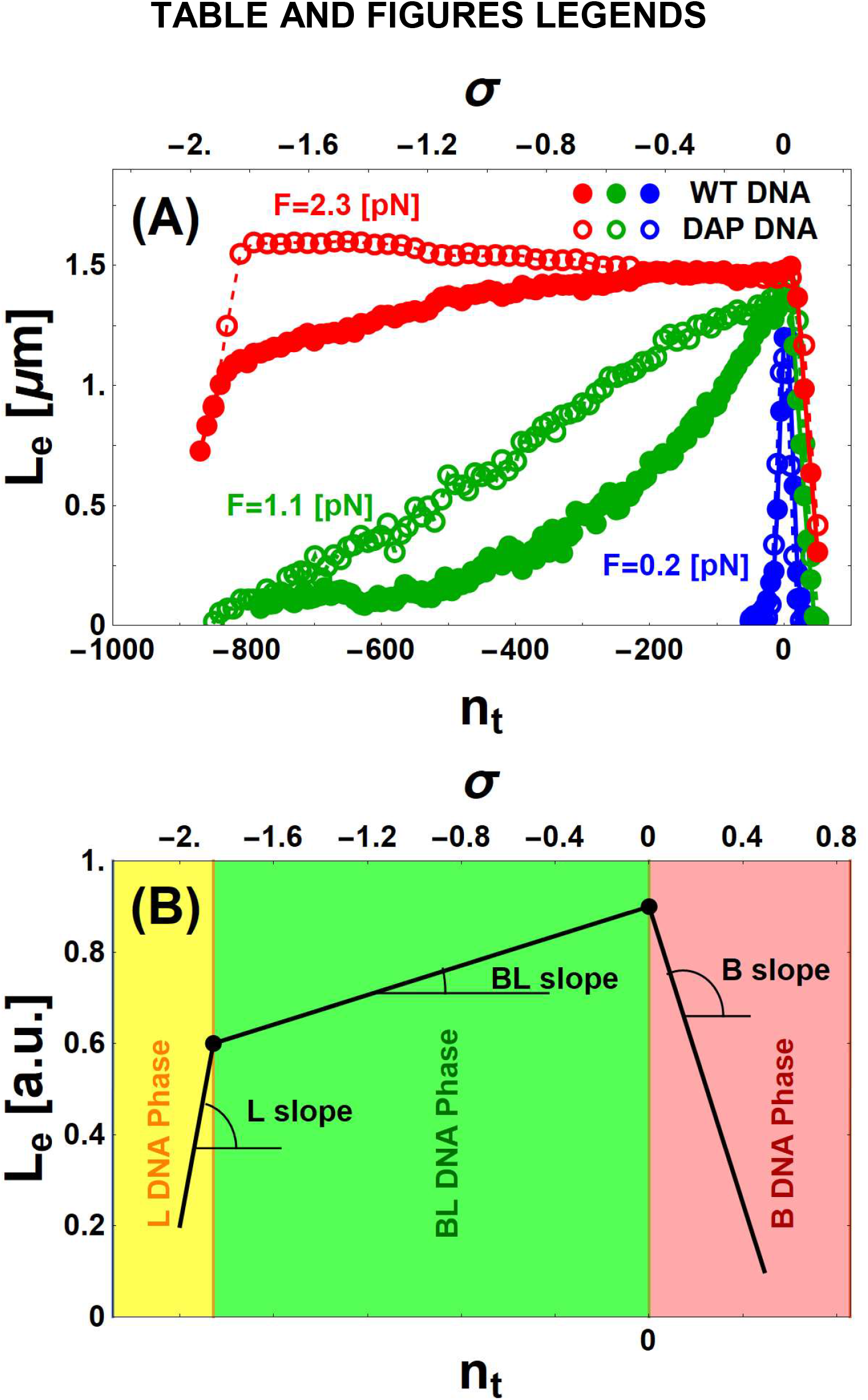
Torsional behavior of WT and DAP DNA. A: Representative MT data for WT (filled disks) and for DAP (open circles) DNA display different extensions, L_e_, as a function of the number of imposed turns n_t_, (lower axis) or supercoiling density *σ*, (upper axis) at fixed values of force F. B: Simplified sketch of theoretical DNA extension, L_e_, obtained as a function of the number of imposed turns n_t_, or supercoiling density *σ*, at fixed values of F shows three phases. The different, color-coded DNA phases are: L (yellow), mixed BL (green), and B (red). The specified angles illustrate the different slopes dL_e_/dn_t_ characterizing the L, BL, and B phases.

These three regimes are highlighted in Figure 1B, a schematic representation of a typical L_e_ vs n_t_ curve which subdivides it into three linear segments, where L_eB_, L_eBL_,and L_eL_ define the DNA extension in the B, BL, and L forms (BL indicates a mixture of B and L forms). Each segment is characterized by a slope, dL_e_/dn_t_. For n_t_>0 (red region), the DNA remains in the B-form and dL_eB_/dn_t_ is negative. For intermediate negative values of n_t_ (green region), the DNA extension can either decrease or increase as a function of n_t_, corresponding to a shallow positive or negative BL slope dL_eBL_/dn_t_, respectively. For higher negative values (<-1.8; yellow region), the DNA is in the L-form and dL_eL_/dn_t_ is positive and steep.

Following these observations, we first adopted a phenomenological approach to study the B, BL and L slopes for WT and DAP DNA, respectively, and for various applied force values.

#### DNA B-form

As shown in Figure 2, the absolute value of the slopes dL_eB_/dn_t_, calculated for the region of the experimental L_e_ vs n_t_ curves where n_t_>0 (B slopes), is a decreasing function of the force. We note that the DAP slope absolute values, despite the large error bars, appear systematically larger than the WT DNA slope absolute values.

**Figure 2.**
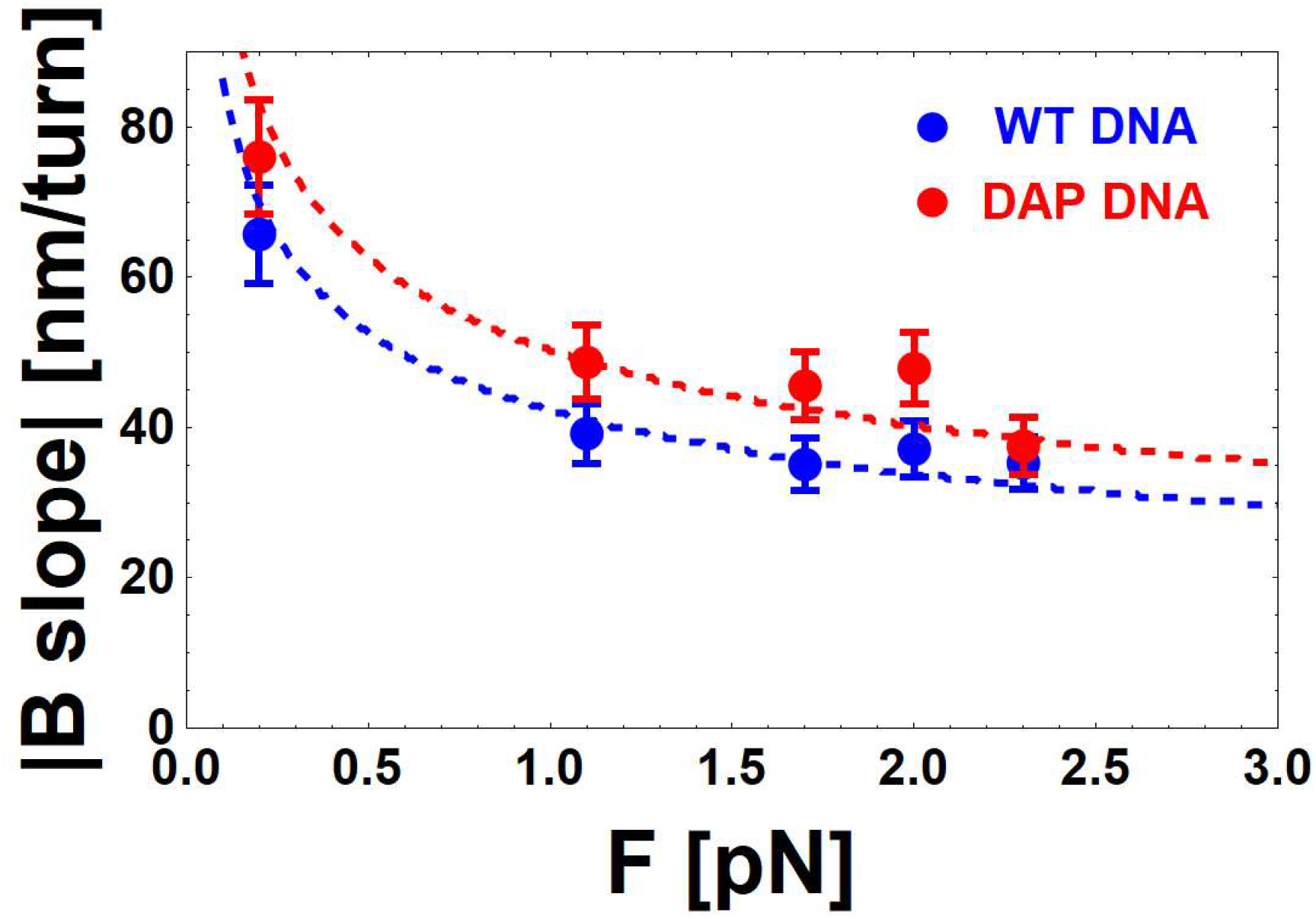
Force dependence of the slope of the extension versus twist data for B-formWT and DAP DNA. Experimental values (dots) and theoretical predictions (dashed curves) for the slope dL_eB_/dn_t_ of B-WT (blue) and B-DAP (red) DNA. Slopes are given in absolute value. Predictions were obtained using the model (22) described in the text assuming: I_s_=150 [mM] and the best fit values L_pB_=40 [nm] (blue); L_pB_=80 [nm] (red).

#### DNA BL-form

The slopes dL_eBL_/dn_t_ (BL slopes), calculated in the intermediate BL region, are shown in Figure 3. The figure reveals that at fixed applied force the slopes of DAP DNA are significantly lower than those of WT DNA. Furthermore, the BL slopes show a change in sign for both DNA types, indicating an inversion force F*, marked by stars in Figure 3, where the BL slope is zero. The inversion force is higher for WT DNA (F*_WT_=2.7 ± 0.3 [pN]) than for DAP DNA (F*_DAP_=1.6 ± 0.3 [pN]).

**Figure 3.**
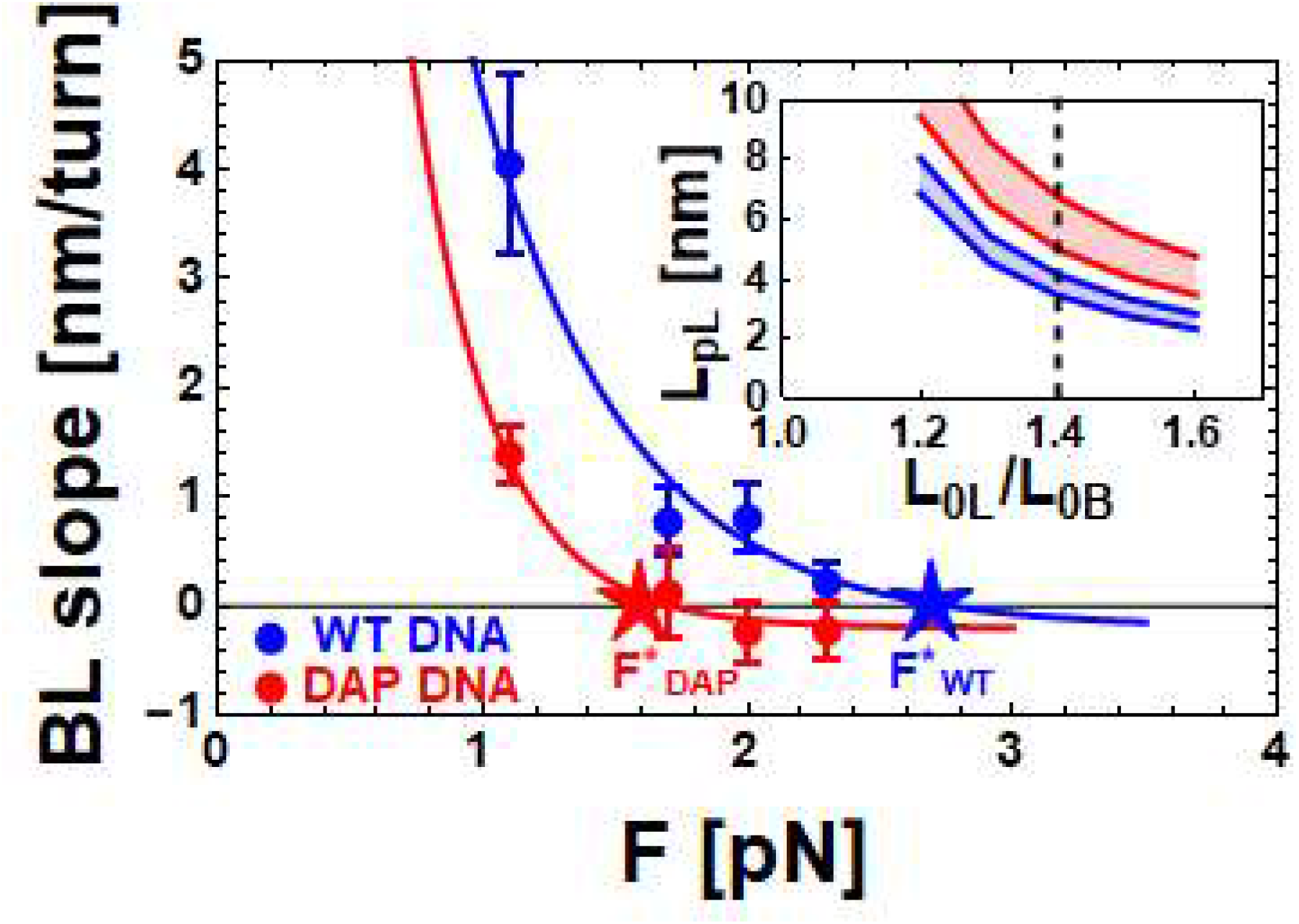
Force dependence of the slope of the BL phase for WT and DAP DNA. Experimental data (dots) and theoretical predictions (solid lines) for the slope dLeBL/dnt of the BL-DNA phase are shown as a function of the applied force F for WT (blue) and DAP (red) DNA. The continuous lines are a guide for the eye. F*_DAP_=1.6 [pN] and F*_WT_=2.7 [pN] (stars), represent the measured inversion force for DAP and WT DNA, respectively. Inset: theoretical predictions of the regions of the values of L_pL_ and L_0L_/L_0B_ compatible with the measured values of the inversion forces (red region for F*_DAP_ ± 0.3 [pN] and blue region for F*_WT_ ± 0.3 [pN], respectively). The vertical dashed line represents the assumed values of L_0L_/L_0B_ for WT and DAP (see text for details).

#### DNA L-form

Finally, Figure 4 shows the dependence of the L slopes dL_eL_/dn_t_ (L slopes) on the applied force for both WT and DAP DNA. The values of L slopes are smaller with respect to the corresponding B slopes. Furthermore the L slopes for DAP appear significantly higher than those for WT DNA, highlighting, also in this case, the difference in the response to torsion between WT and DAP DNA.

**Figure 4.**
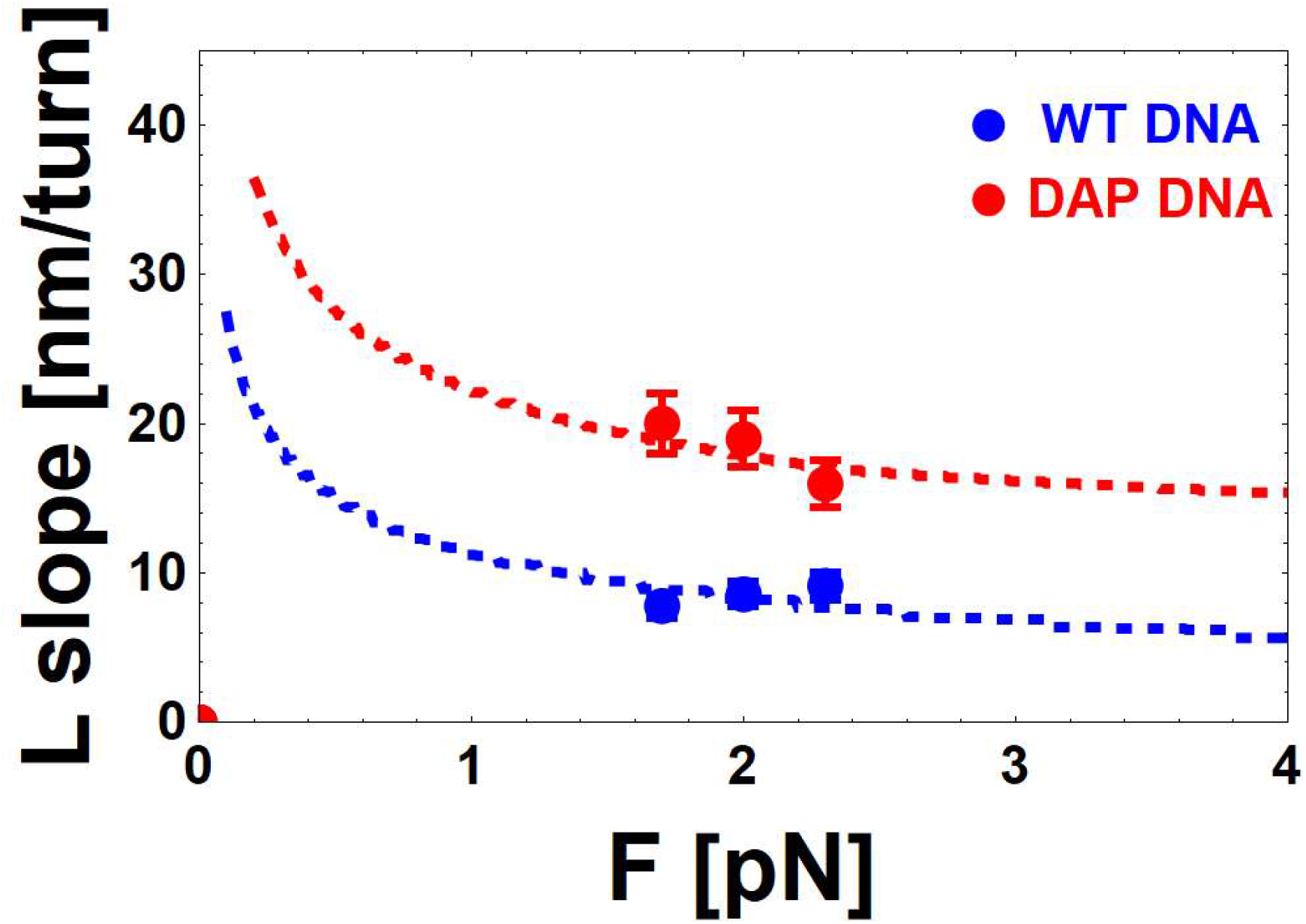
Force dependence of the experimental data (symbols) and the theoretical predictions (dashed lines) for the slopes dL_eL_/dn_t_ of the L-phase for WT (blue) and DAP (red) DNA. Parameters used in the model (22) are as described in the text: ionic strength I_s_=150 [mM], L_pL_=3 [nm] (red line), and L_pL_=1.5 [nm] (blue line) and reduced linear charge density and DNA radius (see text for details).

### Theoretical frame

The variation of the DNA extension under an imposed torsion is due to the need of DNA to relax the torsion by forming plectonema and/or denaturation bubbles (28). For small applied forces (F<0.5 [pN]), the DNA is in B-form and the formation of plectonema is strongly favoured, consequently the variations in DNA extension as a function of torsion are significant. At negative and intermediate values of n_t_ or*σ* (−800<n_t_<-50; −1.8< *σ* <-0.1) and higher values of applied force (F>1 [pN]), i.e. in the DNA BL-form regime, the formation of denaturation bubbles induces slight variations in the DNA extension and the number of base pairs involved in such denaturation bubbles is proportional to n_t_ or *σ* (29). For large n_t_ or *σ*absolute values (n_t_ <-800; *σ*<-1.8) the DNA is completely denatured, its conformation changes from B to L, and any further decrement of n_t_ or likely induces the formation of plectonema in segments of the L-form DNA.

In the following, we propose a simplified theoretical explanation for understanding the torsional behavior of the three DNA forms discussed thus far in either WT, or DAP, DNA.

#### DNA B-form

The variations of the DNA extension under positive imposed turns, or negative turns at low forces, is due to the formation of plectonema (30,31,32). A precise modeling of the geometrical structure of such plectonema has been already proposed, and it successfully predicts quantitatively the B slope of experimentally measured L_e_ vs n_t_ curves in WT DNA (33,34,35,36). Those authors extracted the geometrical conditions of the spiral-like plectonema which minimize the total energy by calculating the total energy as the sum of the potential, bending and electrostatic energies (34). The resulting prediction for the B slope according to that model are the dotted red (DAP) and blue (WT) curves in Figure 2, calculated at 150 [mM] NaCl for a WT DNA of persistence length L_pB_=80 [nm] and L_pB_=40 [nm], respectively. Such specific L_pB_ values are the best fit parameters for the proposed model (34) to the data and are qualitatively similar to the already reported values of persistence length of WT and DAP DNA (5,9) (see Discussion).

#### DNA BL-form

By further decreasing, especially in DNA molecules under increased tension, the DNA extension is well described by a combination of B and L-DNA (mixed BL phase) (33,37). Standard B-DNA is characterized by a persistence length, L_pB_L_0B_=0.34 [nm]. The same parameters are generally difficult to estimate for L-DNA, but a value of L_pL_ [nm] and a corresponding increment of the extension L_0L_/L_0B_= 1.41 have been reported for WT DNA (33). Overall, L-DNA appears to be more flexible and elongated with respect to B-DNA. The percentage of L-DNA induced by torsion increases linearly with*σ*, until the critical value of n_t,max_-800, or*σ* _max_ −1.8, where all B-DNA is converted to L-DNA. At the two extremes (n_t_=0 or*σ* =0 and n_t,max_ −800 or *σ*_max_-1.8), the DNA extension, L_eB_ and L_eL_, respectively, is correctly modelled as a function of the external force F, by a classical Worm Like Chain (WLC) (38,39,40) with the corresponding L_0B_, L_pB_ and L_0L_, L_pL_ for pure B and L-DNA, respectively (41). The relation between F and both L_eB_ and L_eL_ is described by the function *f* given by the classical WLC model as follows:

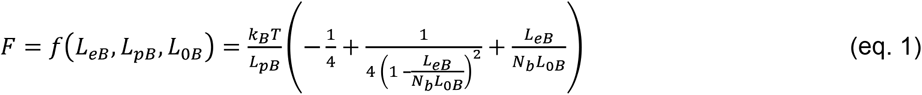

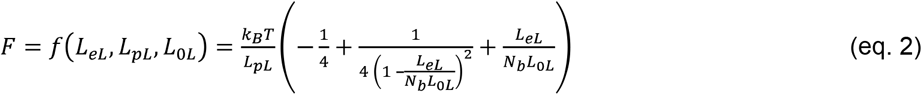

From which, it is possible to calculate the DNA extension as *L*_*eB*_ *= f* ^*-*1^(*F*, L_*pB*,_ L_0*B*_) and L_*eL*_*= f* ^*-*1^(*F*, L_*pL*,_ L_0*L*_).

At intermediate *σ*values the DNA extension L_eBL_ of the BL mixed phase can be described as a linear combination of the WLC contribution from the two pure phases B and L:

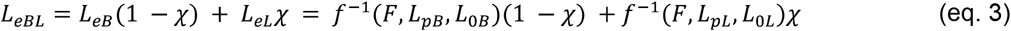

where represents χ the fraction of L-DNA present in the sample and it can be evaluated as

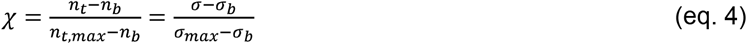

where n_b_ or*σ* _b_ are the number of turns, or supercoiling, corresponding to the buckling transition, the point at which the first supercoil forms, and n_t,max_ or *σ*_max_ are the number of turns, or supercoiling, necessary for a complete conversion to the L-form (*σ* _max_=n_t,max_/(N_b_/10.4)).

The results of this heuristic model are represented in Figure 5A, where the WLC predictions for L_eBL_ *vs* F are shown in the case of the pure B (orange) and L (cyan) phases. The two limit cases of pure B and pure L phases correspond to the values of n_t_=0 and n_t_=n_t,max_, respectively. As apparent from the Figure, the WLC model for the B-form is characterized by both a lower value of the asymptotic extension (L_0B_ < L_0L_) and a higher value of the persistence length with respect to the L-form (L_pB_ > L_pL_). The red star (Figure 5A) indicates the force at which extensions of B and L phases are equivalent, i.e., the inversion force.

**Figure 5.**
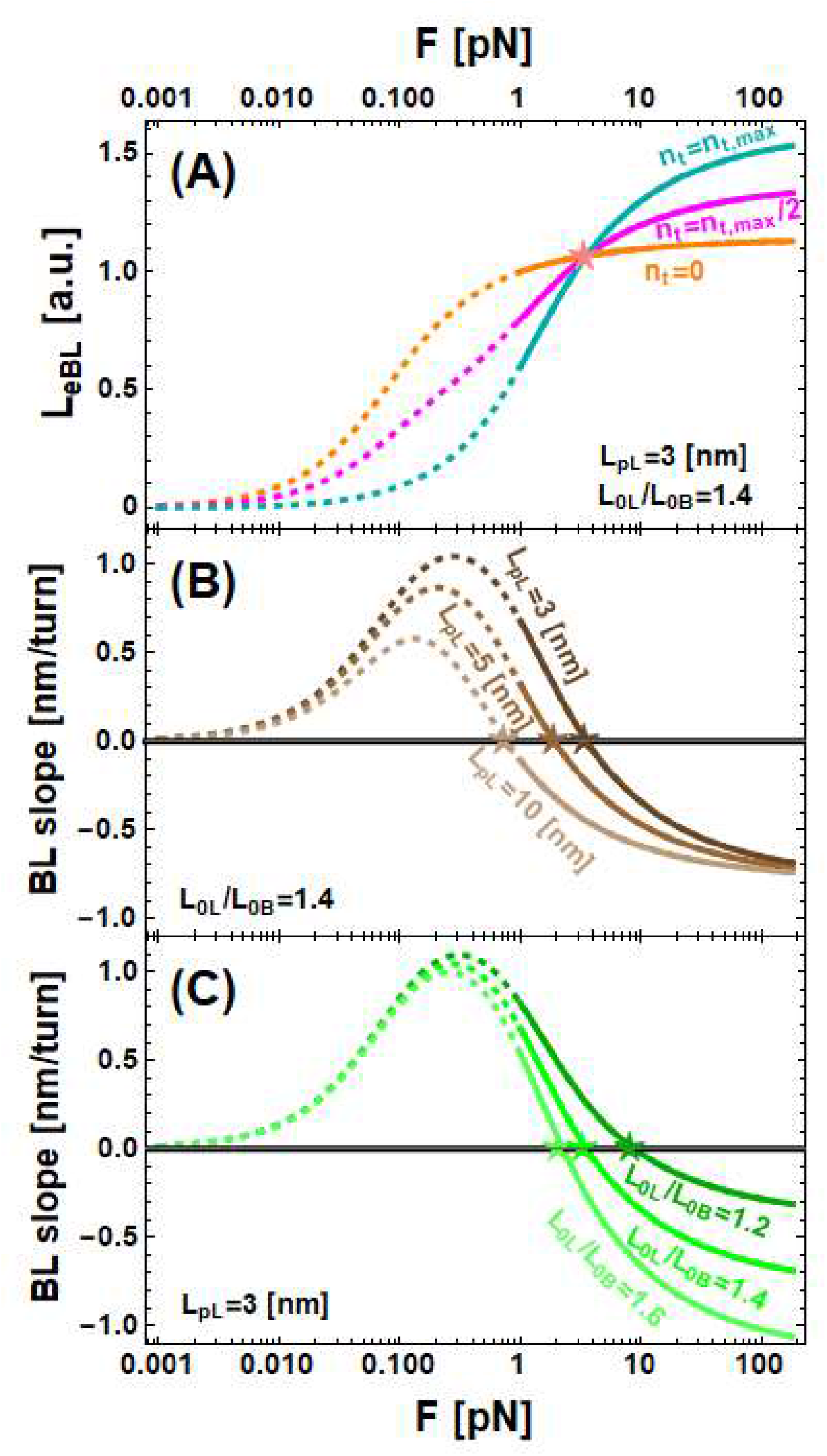
Simulations of DNA extension L_eBL_ and BL-phase slope, dL_eBL_/dn_t_, calculated according to the model described in the text. The theoretical predictions of the model are presented as dashed lines (outside the range of validity) or continuous lines (inside the range of validity). A: The calculated DNA extension L_eBL_ is plotted as a function of the applied force F with fixed persistence length of the L phase, L_pL_=3 [nm], and ratio between the L-form DNA and B-form DNA extension, L_0L_/L_0B_=1.41, for various n_t_ (n_t_=0 (orange); n_t_=n_t,max_/2 (magenta); n_t_=n_t,max_ (cyan)). B: The calculated BL-phase slope is plotted as a function of force with fixed L_0L_/L_0B_=1.41 at L_pL_=3 [nm]; L_pL_=5 [nm]; L_pL_=10 [nm]. C: The calculated BL-phase slope is plotted as a function of force with fixed L_pL_=3 [nm] at L_0L_/L_0B_=1.2; L_0L_/L_0B_=1.4; L_0L_/L_0B_=1.6.

For intermediate values of, this simple model predicts a behavior of L_eBL_ *vs* F described by the magenta curve, obtained for the exemplificative value of n_t_=n_t,max_/2 (Figure 5A). Furthermore, this model allows calculation of the value of the BL slope as dL_eBL_/dn_t_ (Figures 5B and 5C) taking the n_t_ derivative of eq. 3. In all the plots of Figure 5, the theoretical predictions of the model are presented as dashed lines (outside the range of validity) or continuous lines (inside the range of validity), to emphasize that the proposed simplified model is valid only for high force values, where the percentage of denaturation bubbles is higher, and we can disregard the plectonema formation. Given the values of L_0B_ and L_pB_, the free parameters of the model are the persistence length L_pL_ and the extension increment L_0L_ of the L-form. Consequently, in Figure 5B the predictions of the BL slopes are obtained by keeping constant the L_0L_ and considering a variable L_pL_, while viceversa in Figure 5C. The main result of the simplified model is the prediction of the inversion force F* (stars in Figure 5) where the BL slopes assume zero values for different values of L_0L_, and L_pL_. These inversion forces have been consistently calculated for several conditions (Figure S2) where we show F* obtained by keeping constant the L_0L_/L_0B_ ratio and by varying L_pL_ (Figure S2A), and vice versa (Figure S2B). From the predicted inversion force values it is possible to estimate the values of L_0L_/L_0B_ and L_pL_ compatible with the experimentally measured values of F*_WT_=2.7±0.3 [pN] and F*_DAP_=1.6±0.3 [pN] which are indicated by the blue and red continuous horizontal lines in Figure S2 together with their confidence range indicated by horizontal dashed lines. Indeed, by keeping constant the values of L_pB_ and L_0B_, it is possible to calculate a specific value of the inversion force by assuming the values L_pL_ and L_0L_. Vice versa from a specific value of F* it is possible to calculate the values of L_pL_ and L_0L_ compatible with such force values. Namely, the intersections of the horizontal lines of Figure S2 with the calculated lines at various L_0L_/L_0B_ and L_pL_ indicates the L_pL_ and L_0L_/L_0B_ values compatible with the measured values of F*. The results are shown in the inset of Figure 3, where in the space of the two variables L_0L_/L_0B_ and L_pL_, we show the locus of values compatible with the range of measured values of the inversion force, represented as red and blue regions for DAP and WT, respectively.

#### DNA L-form

The L slopes are obtained at high negative values of the imposed supercoiling (n_t_<-800 or *σ*<-1.8). In such a regime, all the DNA is in the L-form and any further torsion presumably induces the formation of plectonema of L-DNA. Since the physical phenomenon of plectonema formation is the same in both cases and the only difference is represented by the specific parameters of the phase under investigation, it would be possible to apply the same model (34) used to explain the B slopes also for L slopes. In Figure 4, we reported the experimentally measured values (symbols) and the values of the L slope predicted by the model (dashed lines) obtained for a specific parameter set: L_pL,DAP_=3 [nm] (red line) and L_pL, WT_=1.5 [nm] (blue line) and reduced values of linear charge density and DNA radius (see Discussion).

## DISCUSSION

The experimental data presented here show significant analogies and differences between WT and DAP DNA. In particular, the measurements of the DNA extension under the effect of controlled mechanical stress, such as imposed torque and stretching, give access to the nanomechanical characterization of WT and DAP DNA in both the B and L forms. Indeed, using MTs it is possible to control the fraction of L-form. Such precise control is hardly achievable outside of the frame of a single molecule experimentation.

### DNA B-form

Firstly we note that, in the regime of positive supercoiling and negative supercoiling at low forces (<0.5 pN), both the WT and DAP-substituted DNA adopt the classical right-handed double helix with unperturbed hydrogen bonds between the Watson and Crick bases (DNA in the B-form). In these regimes, the slope of the dependence of L_e_ *vs*. number of imposed turns, n_t_, is due to the plectonema geometry, which depends on the B-DNA characteristics such as the persistence length and on the tension in the molecule (33,34,35,36,42). In the data presented here (Figure 2) we systematically observe steeper slopes for B-form DAP DNA with respect to WT DNA. The steeper slope of the of L_e_ for DAP DNA *vs*. the number of imposed turnes, n_t_ at low forces, indicates that the persistence length of DAP is definitely larger than that of WT DNA. The prediction of the model we used (34 or https://home.uni-leipzig.de/mbp/index.php/software/) quantifies precisely the B slope dependence on the persistence length, force, and ionic strength. Accordingly, the dashed lines in Figure 2 show the slopes calculated following the model in ref. (34), assuming a best fit value of persistence length L_p_=80 ± 15 [nm] for DAP and 40 ± 10 [nm] for WT B-DNA. The persistence length of DAP-substituted DNA obtained from MT measurements has been already reported in literature (5,9) and is compatible with our best fit values.

#### DNA BL-form

The negative supercoiling region at higher forces (>1pN) exhibits significant differences between the BL and L slopes of WT and DAP-DNA samples. The difference between the inversion force F* of DAP and WT DNA (Figure 3) is striking. This inversion force difference can be explained with the simplified model we obtained by combining in series a WLC model of B- and one of L-form DNA, each with its own characteristic L_p_ and L_0_ parameters.

Based on the assumption that the relative fraction χ of L-DNA is linearly dependent on the number of imposed turns, our model can predict persistence length L_pL_ and contour length L_0L_values, which are compatible with the measured inversion force F*.

Our model is not sufficient to describe in more detail the L-form DNA mechanical parameters. Indeed, in principle, many pairs of values for L_pL_ and L_0L_/L_0B_ within the areas shown in the inset of Figure 3 satisfy the model. Nevertheless, it is possible to determine independently L_0L_/L_0B_ from our data with a simple geometrical reasoning. The argument is based on the observation that all the DNA phases share the same length of the backbone and the differences in the contour lengths are related to the helix geometrical properties. In particular, the DNA contour length is determined by the helical pitch that can be easily calculated from the threshold (n_t_=-800) at which the BL region ends. A simple calculation (see Figure S3 for details) leads to a value L_0L_/L_0B_=1.4 equal for both WT and DAP, since both have the same threshold in n_t_=-800 for the BL to L transition. This value of 1.4 is also reported in the literature only for WT DNA (33,41). By using this value for L_0L_/L_0B_, we obtained the persistence length values L_pL,WT_ ≈3.5[nm] and L_pL,DAP_ ≈7.0[nm] for the L-form.

The resulting increment of the L-DNA persistence length of DAP with respect to WT DNA suggests a more structured and rigid form of the L phase in DAP DNA. This increment of DAP stiffness confirms the qualitative observation that DAP is more easily extended with respect to the WT at constant force (see Figure 1A). Unlike B-DNA, the helically intertwined but non-hybridized L-DNA form is not well defined. The L-form is related to the torsional stress that drives the two strands of DNA counterclockwise while the electrostatic repulsion of the backbone strands gives rigidity to the chain. Some DNA bases might experience large enough negative twist to allow base pairing in the left hand configuration. However, in this case, Z-DNA would result, which is characterized by different mechanical parameters. In particular, for WT Z-DNA has a persistence length, L_pZ,WT_=200 nm, much higher than that of L-DNA (33,43). In the BL region we cannot exclude that a small fraction of Z-DNA phase might be present along the chain. Differing tendencies to generate Z-DNA in the BL region could explain the observed difference of L_pL_determined for DAP DNA.

#### DNA L-form

Finally, the measurements illustrated in Figure 4 shows a steeper dependence of L_eL_ on n_t_ for DAP DNA in the region where all the DNA chain is completely converted to L-DNA. This observation is qualitatively confirmed by the model we adopted (34), since in this region the slope is presumably due to the formation of plectonemes and an increment can be justified with an increase in the persistence length of the L-form. The difference in persistence length qualitatively explains the difference between dL_eB_/dn_t_ and dL_eL_/dn_t_, together with the difference between dL_eL,WT_/dn_t_ and dL_eL,DAP_/dn_t_. Unfortunately, the model used to describe the plectonemes formation for the B-DNA (34) can grasp only qualitatively the behaviour of L-DNA. Indeed, the model fits the data well (see dashed lines of Figure 4) imposing L_pL,WT pL,DAP_ which are different from what we derived above from the analysis of BL slopes (L_pL,WT_ and what reported in the literature (L_pL,WT_ (33). Furthermore, to obtain a reasonable fit with the model (34) is also necessary to impose a reduction of a factor of two of the filament linear charge density and of the DNA diameter, which would result in unrealistic parameters. We note, however, that the differences between the data and the model predictions (34) suggest that the model was extended beyond its range of validity. Assuming that the L slope simply depends on the formation of plectonema is likely naive, since as reported in (33) for the WT DNA, in the L regime, the torque does not seem constant, as should be expected in a plectonemic regime. A possible way to reconcile all these clues is to assume the presence of a new DNA phase after the complete denaturation process. However, whatever the conformation of DNA in this region might be, the large difference between the L slopes attests to a significant difference in the mechanical parameters of DAP with respect to WT DNA in this extremely negatively coiled regime.

In conclusion, L-DNA is the product of enzymes which unwind and/or denature the double helix and a target of regulatory proteins that selectively bind unwound or left-handed DNA. Therefore, given the low levels of unwinding and tension at which it forms, L-DNA likely plays a role in a wide range of DNA transactions. The higher bending rigidity of DAP DNA compared to normal DNA, imposed larger plectonemic gyres at high levels of unwinding. These features emphasize the dependence of the mechanics of L-form DNA on the extent of triple hydrogen bonding and/or on modifications that alter the minor groove and the associated stabilizing spine of water molecules (44) or the major groove in which methylation makes DNA more susceptible to B-Z transitions (45). More generally, the different mechanical properties of DAP expand the tool kit for the rational design of biomaterials.

## DATA AVAILABILITY

Not applicable.

## SUPPLEMENTARY DATA

Supplementary Data are available at NAR online.

## ACKNOWLEDGEMENT

We thank C.A. Marrano for help with the measurements and R. Seidel for discussions about the theoretical model.

## FUNDING

Funding for open access charge: shared between University of Milano-Bicocca and National Institutes of Health [R01GM084070].

National Institutes of Health [R01GM084070 to L.F.].

## SUPPLEMENTARY DATA

**Figure S1.**
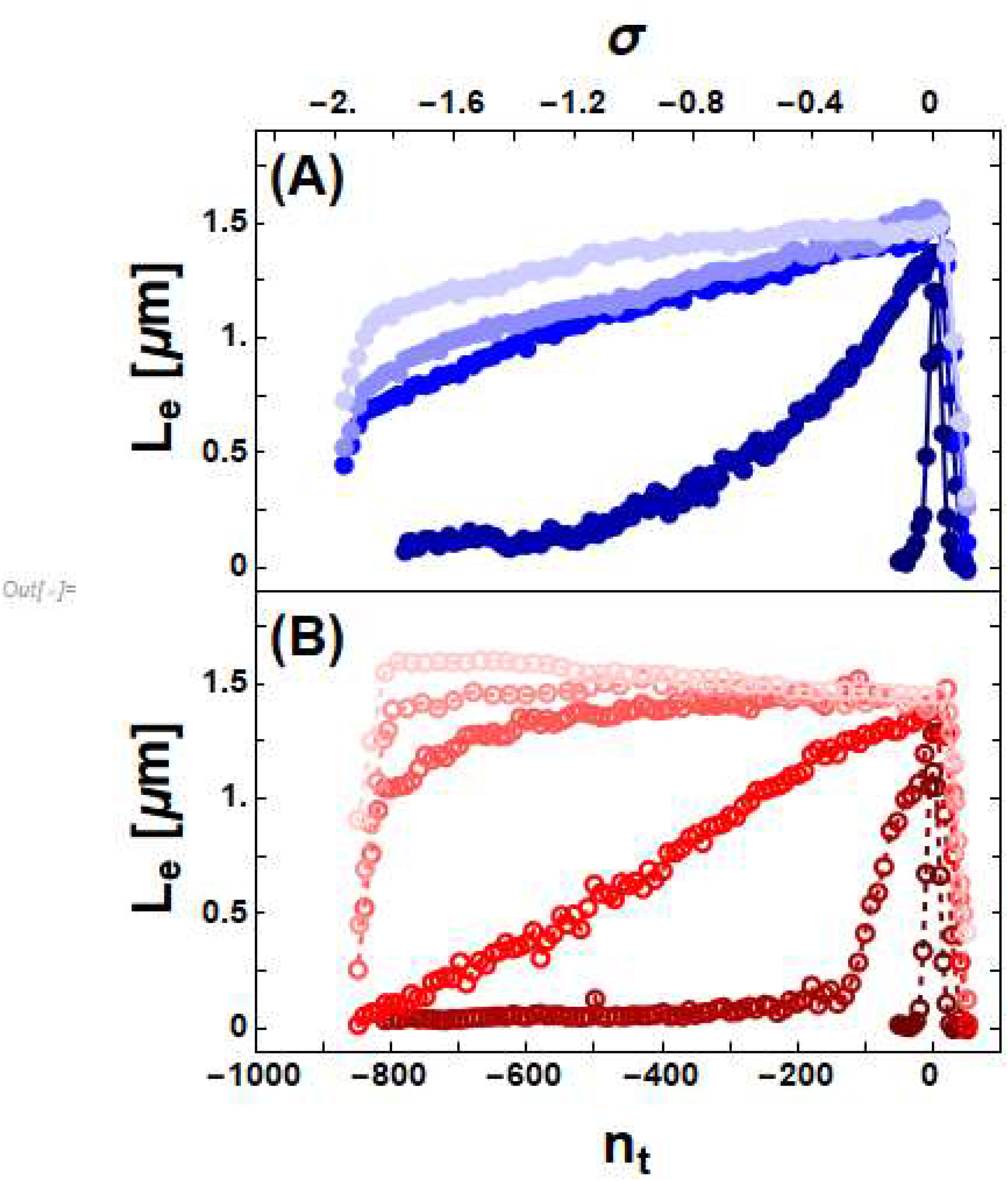
Torsional behavior of WT and DAP DNA. The measured DNA extension, L_e_, is plotted as a function of the number of imposed turns, n_t_, or equivalently supercoiling density, *σ*, for WT (A, filled circles) and DAP DNA (B, open circles) at tensions of 0.2, 1.1, 1.7, 2.0, 2.3 [pN] (light to dark shades of blue or red).

**Figure S2.**
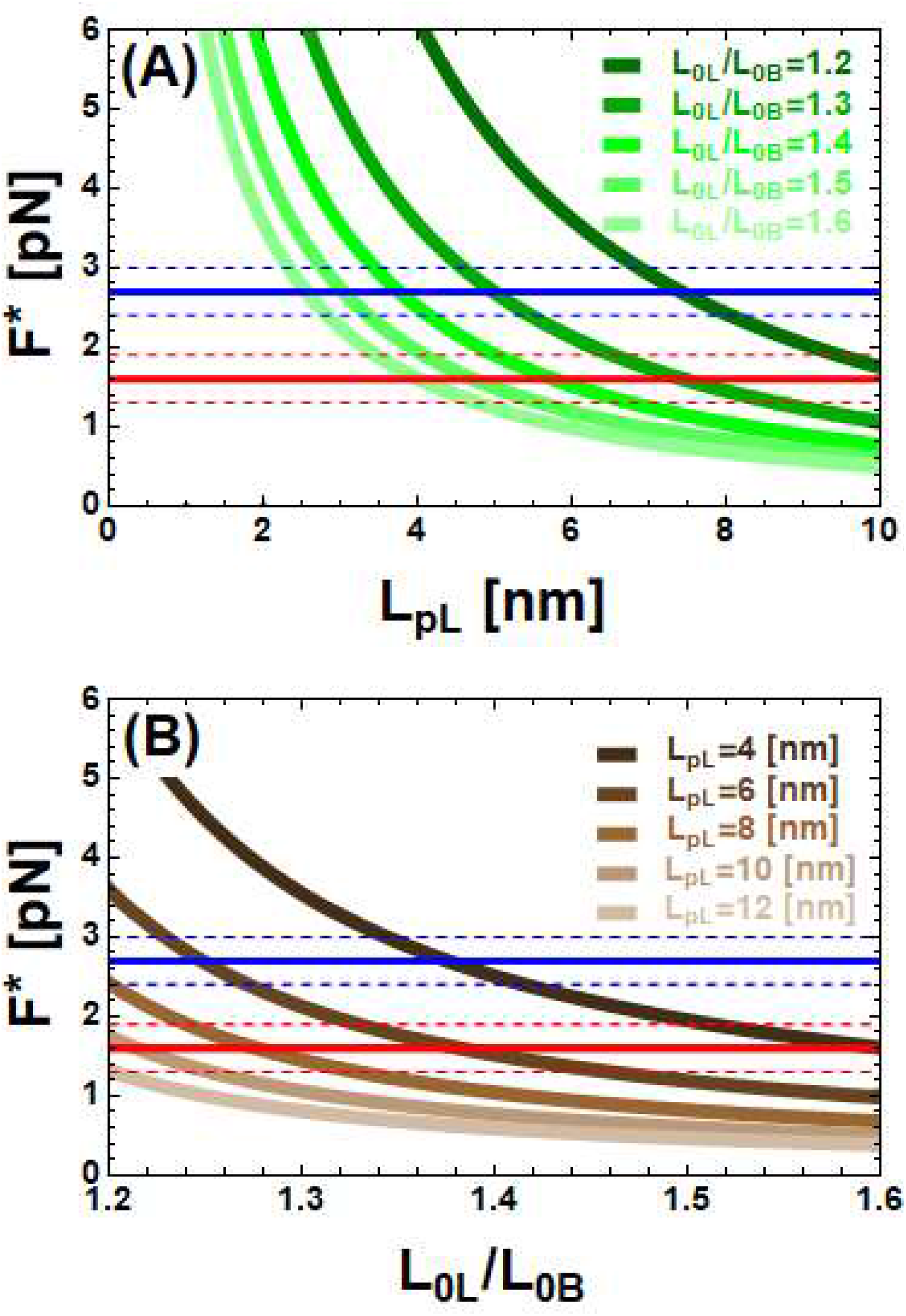
Theoretical values of the inversion force F* calculated as a function of the persistence length L_pL_ of the L phase and ratio L_0L_/L_0B_ between the DNA L and B extension. The continuous horizontal lines represent the measured values of the inversion forces F* for WT (blue line) and DAP (red line) DNA. The dashed lines show the uncertainty of the measured forces (F*_DAP_=1.6 ± 0.3 [pN] and F*_WT_=2.7 ± 0.3 [pN]). F* was calculated as a function of L_pL_ for L_0L_/L_0B_=1.2, 1.3, 1.4, 1.5, 1.6, (A) or as a function of L_0L_/L_0B_ with L_pL_=4, 6, 8, 10, 12 [nm] (B).

**Figure S3.**
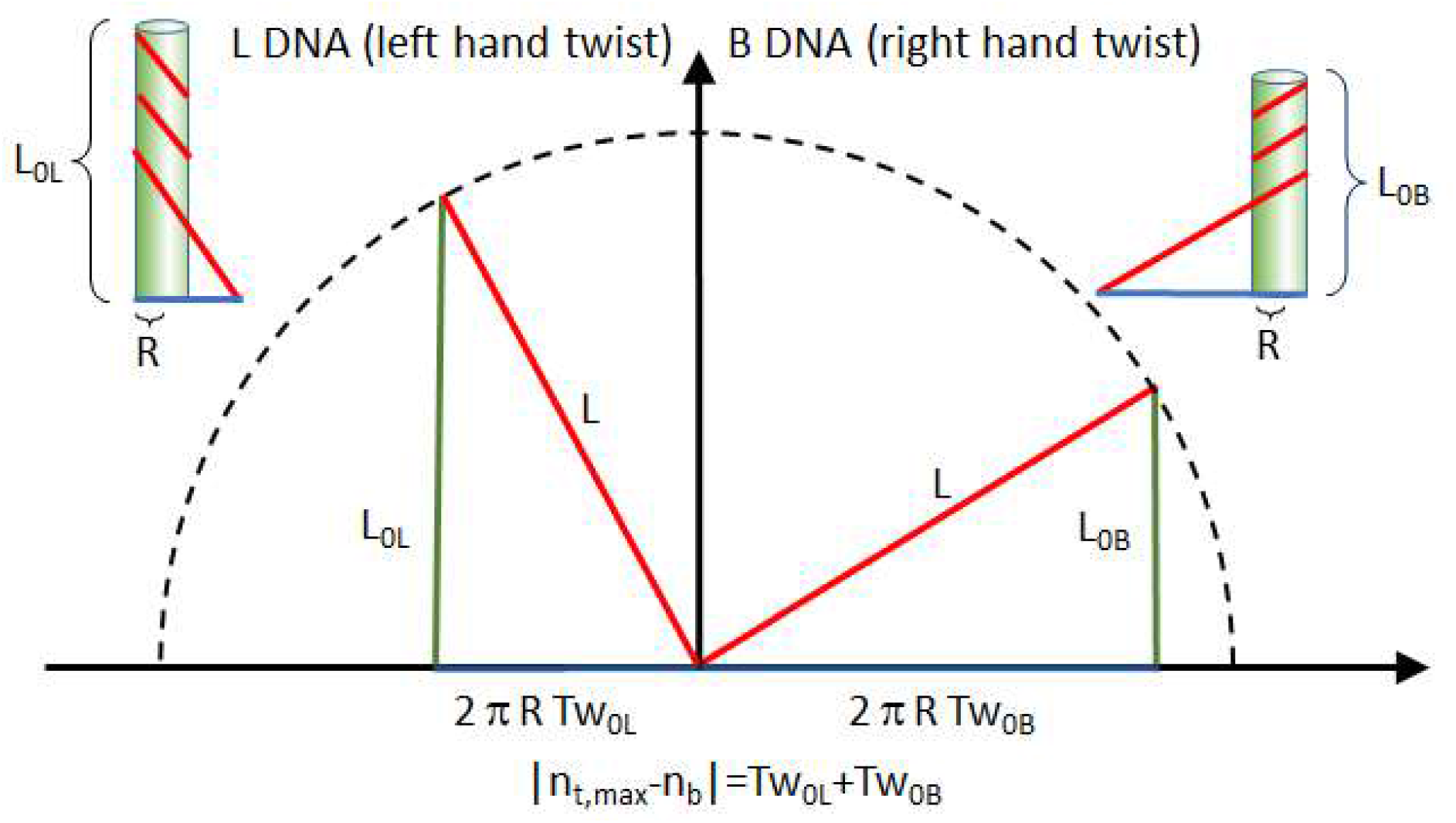
A geometrical model for calculating the contour length increment from B- to L-form DNA. Unwrapping a double-stranded filament, we obtain a triangle (top right for B DNA) and top left for L DNA). The hypotenuse (in red) corresponds to the length of the totally unwrapped filament L=0.6 [nm] per bp (indicated here as the length of one of the two single strands) which is constant and fixes the locus of the points to which the triangle corner is allowed to belong (dashed circumference). The vertical cathetus (in green) of the triangle corresponds to the contour length of the double stranded DNA, L_0L_ or L_0L_. The horizontal cathetus (in blue) is determined by the Pythagorean Theorem, and it corresponds to the circumference of the DNA (2 is the DNA radius) multiplied the twist of the DNA in the different forms Tw_0B_=1 turn/10.4 bps and Tw_0L_, the twist of the DNA form determines its contour length. The relation |n_t,max_-n_b_|=Tw_0L_+Tw_0B_ imposes the topological constraint on the twist number: the total twist applied to the DNA (n_t,max_) is absorbed as twist by the chain (n_b_) which converts the natural twist of the molecule from Tw_0B_ to Tw_0L_. It follows that since WT and DAP DNA share the same threshold n_t,max_, they also share the same Tw_0L_ and, consequently, the same contour length of the L phase for DAP and WT DNA (L_0L,WT_=L_0L,DAP_).

## REFERENCES

1. Peters, J.P., Yelgaonkar, S.P., Srivatsan, S.G., Tor, Y. and Maher, L.J. (2013) Mechanical properties of DNA-like polymers. Nucleic Acids Res., 41, 10593–10604.

2. Bailly, C., Waring, M.J. and Travers, A.A. (1995) Effects of base substitutions on the binding of a dna-bending protein. J. Mol. Bio., 253, 1–7.

3. Khudyakov, I.Y., Kirnos, M.D., Alexandrushkina, N.I. and Vanyushin, B.F. (1978) Cyanophage S-2L contains DNA with 2,6-diaminopurine substituted for adenine. Virology, 88, 8–18.

4. Bailly, C. and Waring, M.J. (2001) Use of DNA molecules substituted with unnatural nucleotides to probe specific drug-DNA interactions. Methods Enzymol., 340, 485–502.

5. Fernández-Sierra, M., Shao, Q., Fountain C., Finzi L. and Dunlap D. (2015) E. coli gyrase fails to negatively supercoil diaminopurine-substituted DNA. J. Mol. Biol., 427, 2305 2318.

6. Howard, F.B. and Miles, H.T., (1984) 2NH2A.T helices in the ribopolynucleotide and deoxypolynucleotide series -structural and energetic consequences of 2NH2A substitution. Biochemistry, 23, 6723–6732.

7. Hoheisel, J.D. and Lehrach, H. (1990) Quantitative measurements on the duplex stability of 2,6-diaminopurine and 5-chloro-uracil nucleotides using enzymatically synthesized oligomers. FEBS Lett., 274, 103–106.

8. SantaLucia, J. and Hicks, D. (2004) The thermodynamics of DNA structural motifs, Annu. Rev. Biophys. Biomolec. Struct., 33, 415–440.

9. Cristofalo, M., Kovari, D., Corti, R., Salerno, D., Cassina, V., Dunlap, D., Mantegazza, F. (2019) Nanomechanics of diaminopurine-substituted DNA. Biophys. J., 116, 760–771.

10. Bailly, C. and Waring, M.J., (1998) The use of diaminopurine to investigate structural properties of nucleic acids and molecular recognition between ligands and DNA. Nucleic Acids Res., 26, 4309–4314.

11. Virstedt, J., Berge, T., Henderson, R.M., Waring, M.J. and Travers, A.A. (2004) The influence of DNA stiffness upon nucleosome formation. J. Struct. Biol., 148, 66–85.

12. Peters, J.P., Mogil, L.S., McCauley, M.J., Williams, M.C., Maher, L.J. (2014) Mechanical properties of base-modified DNA are not strictly determined by base stacking or electrostatic interactions. Biophys. J., 107, 448–459

13. Vorlickova, M., Sagi, J., Szabolcs, A., Szemzo, A., Otvos, L. and Kypr, J., (1998) Conformation of the synthetic DNA poly(amino2dA-dT) duplex in high-salt and aqueous alcohol-solutions. Nucleic Acids Res., 16, 279–289.

14. Saenger, W. (1984) DNA as target molecule for drugs and action of the antimetabolite 6-azauridine, Acta Crystallogr. Sect. A, 4, C58–C58.

15. Vorlickova, M., Kejnovska, I., Bednarova, K., Renciuk, D. and Kypr, J. (2012) Circular Dichroism Spectroscopy of DNA: From Duplexes to Quadruplexes. Chirality, 24, 691–698.

16. Lee, M., Kim, S.H., Hong, S.-C. (2010) Minute negative superhelicity is sufficient to induce the B-Z transition in the presence of low tension. Proc. Natl. Acad. Sci. USA, 107, 4985–4990.

17. Neuman, K.C., Nagy, A. (2008) Single-molecule force spectroscopy: optical tweezers, magnetic tweezers and atomic force microscopy. Nat. Methods., 5, 491–505.

18. De Vlaminck, I., Dekker, C. (2012) Recent advances in magnetic tweezers. Annu. Rev. Biophys., 41, 453–472.

19. Dulin, D., Lipfert, J., Moolman, M.C., Dekker, N.H. (2013) Studying genomic processes at the single-molecule level: introducing the tools and applications. Nat. Rev. Genet., 14, 9–22.

20. Galburt, E.A., Tomko, E.J., Stump, W.T., Ruiz Manzano, A. (2014) Force-dependent melting of supercoiled DNA at thermophilic temperatures. Biophys. Chem., 187, 23–28.

21. Abels, J., Moreno-Herrero, F., van der Heijden, T., Dekker, C., Dekker, N., Single-molecule measurements of the persistence length of double-stranded RNA, Biophys. J., 88 (2005) 2737–2744.

22. Salerno, D., Brogioli, D., Cassina, V., Turchi, D., Beretta, G.L., Seruggia, D., Ziano, R., Zunino, F. and Mantegazza, F. (2010) Magnetic tweezers measurements of the nanomechanical properties of the DNA in the presence of drugs. Nucleic Acids Res., 38, 7089–7099.

23. Tempestini, A., Cassina, V., Brogioli, D., Ziano, R., Giovannoni, R., Cerrito, M.G., Salerno, D. and Mantegazza, F. (2013) Magnetic tweezers measurements of the nanomechanical stability of DNA against denaturation at various conditions of pH and ionic strength. Nucleic Acids Res., 41, 2009–2019.

24. Salerno, D., Beretta, G., Zanchetta, G., Brioschi, S., Cristofalo, M., Missana, N., Nardo, L., Cassina, V., Tempestini, A., Giovannoni, R., Cerrito, M.G., Zaffaroni, N., Bellini, T. and Mantegazza, F. (2016) Platinum-based drugs and DNA interactions studied by single-molecule and bulk measurements. Biophys. J., 110, 2151–2161.

25. Gosse, C. and Croquette, V. (2002) Magnetic tweezers: Micromanipulation and force measurement at the molecular level. Biophys. J., 82, 3314–3329.

26. Allemand, J.F., Bensimon, D., Jullien, L., Bensimon, A. and Croquette, V. (1997) pH-dependent specific binding and combing of DNA. Biophys. J., 73, 2064 2070.

27. Strick, T.R., Allemand, J.F., Bensimon, D. and Croquette, V. (1998) Behavior of supercoiled DNA. Biophys. J., 74, 2016 2028.

28. Allemand, J.F., Bensimon, D., Lavery, R., Croquette, V. (1998) Stretched and overwound DNA forms a Pauling-like structure with exposed bases. Proc. Natl. Acad. Sci. USA., 95, 14152–14157.

29. Strick, T.R., Allemand, J.F., Croquette, V., Bensimon, D. (2000) Twisting and stretching single DNA molecules. Prog. Biophys. Mol. Biol., 74, 115–140.

30. Strick, T.R., Dessinges, M.N., Charvin, G., Dekker, N.H., Allemand, J.F., Bensimon, D., Croquette, V. (2003) Stretching of macromolecules and proteins. Rep. Prog. Phys., 66, 1–45.

31. Fu, W.-B., Wang, X.-L., Zhang, X.-H., Ran, S.-Y., Yan, J., Li, M., (2006) Compaction dynamics of single DNA molecules under tension, J. Am. Chem. Soc., 128, 15040–15041.

32. Lipfert, J., Klijnhout, S., Dekker, N.H. (2010) Torsional sensing of small-molecule binding using magnetic tweezer. Nucleic Acids Res., 38, 7122 7132.

33. Sheinin, M.Y., Forth, S., Marko, J.F. and Wang, M.D. (2011) Underwound DNA under tension: structure, elasticity, and sequence-dependent behaviors. Phys. Rev. Lett., 107, 108102.

34. Maffeo, C., Schopflin, R., Brutzer, H., Stehr, R., Aksimentiev, A., Wedemann, G. and Seidel, R. (2010) DNA-DNA interactions in tight supercoils are described by a small effective charge density. Phys. Rev. Lett., 105, 158101.

35. Neukirch, S. and Marko, J.F. (2011) Analytical description of extension, torque, and supercoiling radius of a stretched twisted DNA. Phys. Rev. Lett., 106, 138104.

36. Lam, P.-M. and Zhen, Y. (2015) Extension, torque, and supercoiling in single, stretched, and twisted DNA molecules. J. Chem. Phys., 143, 174901.

37. R. Vlijm, A. Mashaghi, S. Bernard, M. Modesti, C. Dekker (2015) Experimental phase diagram of negatively supercoiled DNA measured by magnetic tweezers and fluorescence. Nanoscale, 7, 3205–3216.

38. Marko, J.F., Siggia, E.D. (1995) Stretching DNA, Macromolecules, 28, 8759 8770.

39. Bouchiat, C., Wang, M.D., Allemand, J., Strick, T., Block, S.M., Croquette, V. (1999) Estimating the persistence length of a worm-like chain molecule from force-extension measurements. Biophys. J., 76, 409–413.

40. Wang, Y., van Merwyk, L., Toensing, K., Walhorn, V., Anselmetti, D., Fernàndez-Busquets, X. (2017) Biophysical characterization of the association of histones with single-stranded DNA. Biochim. Biophys. Acta-Gen. Subj., 1861, 2739–2749.

41. Marko, J.F. and Neukirch, S. (2013) Global force-torque phase diagram for the DNA double helix: structural transitions, triple points, and collapsed plectonemes. Phys. Rev. E. Stat. Nonlin. Soft Matter Phys., 88, e062722.

42. Mosconi, F., Allemand, J.F., Bensimon, D. and Croquette, V. (2009) Phys. Rev. Lett., 102, 078301.

43. Thomas, T.J. and Bloomfield, V.A. (1983) Chain flexibility and hydrodynamics of the B-form and Z-form of poly(dG-dC).poly(dG-dC). Nucleic Acids Res., 11, 1919–1930.

44. McDermott, M.L., Vanselous, H., Corcelli, S.A., Petersen, P.B. (2017) Hydration. ACS Central Science, 3, 708–714.

45. Behe, M., Felsenfeld, G. (1981) Effects of Methylation on a Synthetic Polynucleotide: The B-Z Transition in Poly(dG-m5dC):poly(dG-m5dC). Proc. Natl. Acad. Sci. USA., 78, 1619–1623.

